# WNT and inflammatory signaling distinguish human Fallopian tube epithelial cell populations

**DOI:** 10.1101/2020.01.15.908293

**Authors:** Ian Rose, Mallikarjun Bidarimath, Alex Webster, Andrew K. Godwin, Andrea Flesken-Nikitin, Alexander Yu. Nikitin

**Author notes:** Correspondence to: Andrea Flesken-Nikitin, or Alexander Yu. Nikitin.

## Abstract

A substantial fraction of ovarian/extra-uterine high-grade serous carcinomas (HGSCs) likely originate in the distal region of the Fallopian tube’s epithelium (TE) before implanting/metastasizing to the ovary. Unfortunately, molecular and cellular mechanisms rendering preferential cancer susceptibility of the human distal TE remain insufficiently elucidated, largely due to limited primary human TE gene expression data. Here we report an in depth bioinformatic characterization of 34 primary TE cell mRNA-seq samples. These samples were prepared from the proximal and distal TE regions of 12 normal Fallopian tubes. TE cells were segregated based on their aldehyde dehydrogenase (ALDH) activity. As compared to the proximal TE, cells from the distal region form organoids with higher frequency and larger size during serial organoid formation assays. Consistent with enrichment for organoid-forming stem/progenitor cells, ALDH+ cells have greater WNT signaling activity. Comparative evaluation of proximal and distal TE cell population’s shows heightened inflammatory signaling in distal differentiated (ALDH-) TE. Furthermore, comparisons of proximal and distal TE cell populations finds that the distal TE express gene sets characteristic of four HGSC molecular sub-types, and that distal ALDH+ cell populations exhibit greater enrichment than their ALDH-counterparts. Taken together, our study shows that increased organoid forming capacity, WNT and inflammatory signaling, and HGSC signatures underlie the differences between distal and proximal regions of the human TE. These findings provide the basis for further mechanistic studies of distal TE susceptibility to the malignant transformation.

## Introduction

Ovarian/extra-uterine high-grade serous carcinoma (HGSC) is the most common and most lethal gynecological malignancy, accounting for nearly 14,000 deaths per year in the United States (Siegel et al., 2020). There is mounting evidence indicating that serous tubal intraepithelial carcinomas (STICs) are precursor lesions to some HGSCs (Kim et al., 2018; Labidi-Galy et al., 2017; Soong et al., 2018). Moreover, in vivo perturbations of TP53, MYC, and hTERT and RB family genes, which are associated with pathways frequently perturbed in HGSC (Siegel et al., 2020), cause human TE cells to adopt traits reminiscent of STICs/HGSCs (Yamamoto et al., 2016). It has been noted that STICs tend to occur more frequently in the distal region (closer to the ovary) than in the proximal region (farther from the ovary) of the Fallopian tube, also known as the uterine tube (Kim et al., 2018; Lee et al., 2007; Nassar and Blanpain, 2016).

Stem cells are frequently implicated in malignant transformation (Flesken-Nikitin et al., 2013; Nassar and Blanpain, 2016), thus regional differences in the TE stem cells may account for the distal TE’s tendency to harbor STICs. Indeed, previous studies based on immunohistochemical analysis of TE sections have suggested that stem/progenitor cells may occur more frequently in the distal TE (Paik et al., 2012; Schmoeckel et al., 2017). Support for this notion also comes from the observations of preferential sphere formation by human distal TE (Paik et al., 2012) and organoid formation of the mouse distal oviduct (the mouse analogue to the Fallopian tube (Xie et al., 2018). Long-term organoid formation assays have been indicative of adult tissue stem cells being present in human TE cells isolated from both the proximal and distal regions of the Fallopian tube (Kessler et al., 2015). However, quantitative comparisons of proximal and distal TE organoid capacity have not been performed. Additionally, studies which interrogate organoid-forming cells, performing quantitative organoid assays and measure global gene expression data in primary human TE are sparse or absent.

Differences between TE stem cell populations are not the only factors that may promote malignant transformation in the distal TE. Chronic inflammation is known to cause cancer in a number of contexts (Elinav et al., 2013). The ovary is known to release inflammatory factors on a regular basis in humans ((Lewandowska et al., 2019; Zamah et al., 2015) and Supplementary Figure 1). The expression of pro-inflammatory cytokine IL-8 has been shown to correlate with ovulation (Palter et al., 2001). Consequently, the ovary-derived factors have long been suspected of promoting malignant transformation (Fathalla, 1971). More recent studies find that follicular fluid induces DNA damage and proliferation in TE (Brachova et al., 2017; King et al., 2011), and exposure to follicular fluid also induces changes in TE reminiscent of STICs (Bahar-Shany et al., 2014). Gene expression data from different human TE cell populations may aid in determining the immediate relevance of these observations to the human TE.

Given that information pertaining to TE stem and differentiated cells is sparse, and gene expression data in primary human cells is very limited, we devised a fluorescence activated cell sorting (FACS) strategy based on ALDH activity to purify populations of stem/progenitor and differentiated epithelial enriched cell populations from both the proximal and distal regions of the human TE. Aldehyde dehydrogenase (ALDH) is a detoxifying enzyme and its increased activity is frequently observed in stem/progenitor cells of ovarian surface epithelium, mammary, prostate, colon, haematopoietic, neural and mesenchymal cell lineage (Douville et al., 2009; Flesken-Nikitin et al., 2013; Ma and Allan, 2010). Long term organoid formation assays demonstrate that ALDH+ cell populations have a greater capacity for organoid formation than ALDH- cell populations. Based on a thorough bioinformatic characterization of the isolated cell populations we also report that ALDH+ cells have greater WNT signaling activity and that the distal TE is characterized by increased inflammatory signaling and gene expression patterns reminiscent of HGSC.

## Results

### ALDH activity distinguishes organoid-forming cells

All Fallopian tubes used in our experiments were removed from donors not afflicted with ovarian cancer and not carrying mutant BRCA1/2 alleles, and who were between the ages of 32 and 51. Proximal and distal regions of the TE were divided as indicated in Figure 1A. To test if there are regional differences in TE organoid formation, we prepared organoids from distal and proximal regions of the Fallopian tubes and propagated them for 4 passages. Consistent with previous report (Kessler et al., 2015), primary TE cells from both distal and proximal regions were able to form organoids (Figure 1B). However, distal TE cells consistently formed organoids at a significantly higher frequency than their proximal region counterparts (Figure 1C). Furthermore, organoids grown from the distal TE region tend to be significantly larger than their proximal counterparts (Figure 1D). Both distal and proximal organoids contained ciliated (AcTub+) and secretory (PAX8+) cells, as well as cells expressing stem/progenitor cell marker ALDH1A1.

**Figure 1.**
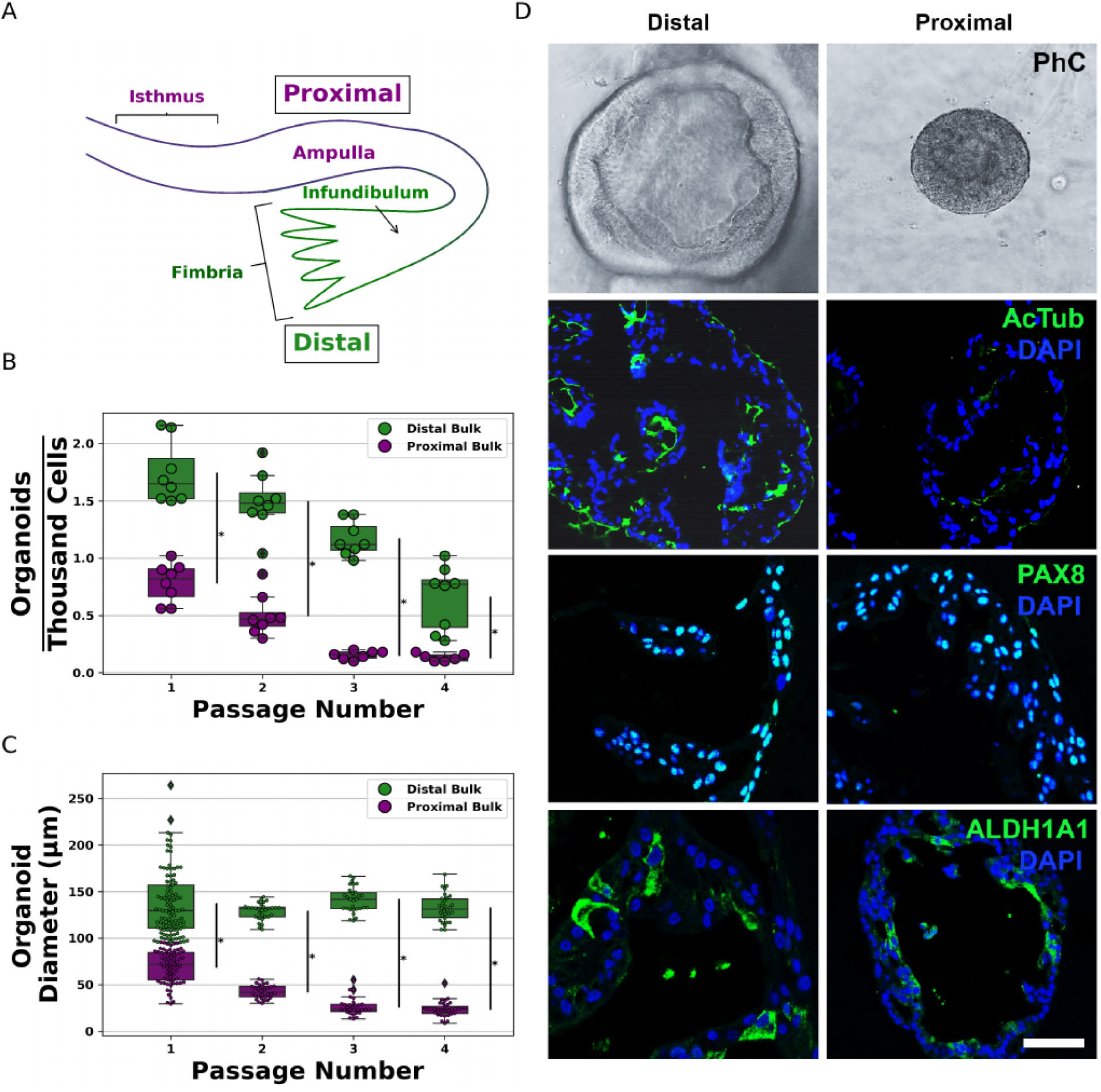
Bulk Fallopian tube organoid formation assays. (A) Schematic of the proximal and distal regions of Fallopian tubes. The frequency (B) and the diameter (C) of the distal and proximal TE orgnoids were measured on the 8th day in the culture successively for 4 passages. Each dot corresponds to an observation. (D) Representative images of human distal and proximal tube epithelial organoids. First row shows the phase contrast (PhC) images of the organoids. The expression of ACTUB (second row), PAX8 (third row), and ALDH1A1 (fourth row) in distal and proximal tubal epithelial organoids. Note the cilia are marked by ACTUB (green), nuclear staining (green) for PAX8 and cytoplasmic staining (green) for ALDH1A1. Counterstaining with DAPI for immunofluorescence staining. Bar, D, First row, 150 μm; Second row, 60 μm; third row, 50 μm; and fourth row, 50 μm. All error bars denote s.d.

Based on previous observations that ALDH activity is frequently observed in stem/progenitor cells, we hypothesized that ALDH+ epithelial (EpCAM+) cell populations have increased organoid formation as compared to ALDH-/EpCAM+ cells. Therefore, we FACS isolated viable EpCAM+/ALDH+ and EpCAM+/ALDH- cell populations from the proximal and distal regions of Fallopian tubes and conducted organoid formation assays (Figure 2A). Light scatter gating was used to exclude debris (Supplementary Figure 2) before collecting viable (Figure 2B) EpCAM+/ALDH+ and EpCAM+/ALDH- cells from proximal and distal regions of the TE (Figure 2C). We have found that both proximal and distal TE EpCAM+/ALDH+ and EpCAM+/ALDH- cell populations have the capacity to form organoids and that significantly more organoids have formed from EpCAM+/ALDH+ isolates, as compared to EpCAM+/ALDH- cell population, in both proximal and distal samples (Figure 2D). Organoid formation is generally indicative of stem/progenitor cells ex vivo. Thus, these findings suggest that ALDH activity is a suitable means of enriching for stem/progenitor cells in human TE isolates.

**Figure 2.**
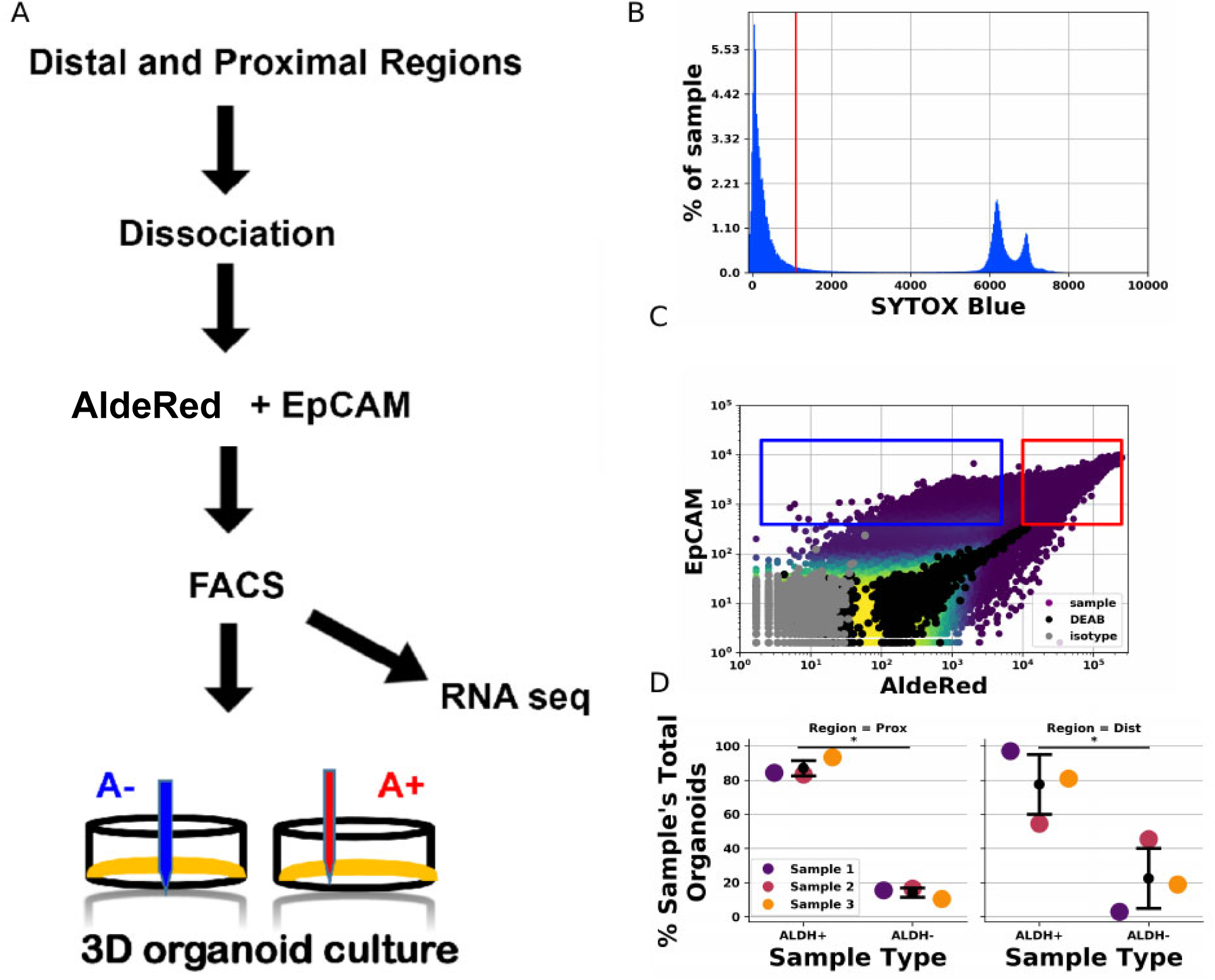
Organoid assays from Fallopian tube epithelial cells isolated by ALDH activity. (A) Schematic describing experimental strategy. Epithelial cells from cryopreserved proximal and distal region Fallopian tube fragments were isolated by FACS on the basis of ALDH activity. (B-C) Representative FACS data. (B) SYTOX Blue staining was used to identify non-viable cells. Cells with SYTOX Blue staining lower than indicated by the red line were declared viable. (C) Staining of EpCAM isotype controls (gray dots) was used to identify EpCAM+ cells. AldeRed staining of cells treated with ALDH inhibitor DEAB (black dots) was used to identify ALDH+ cells. Blue and Red rectangles indicate EpCAM+/ALDH- and EpCAM+/ALDH+ populations, respectively. (D) Organoid formation by EpCAM+ ALDH+ and EpCAM+ ALDH- cells isolated from proximal and distal regions of Fallopian tubes.

### Proximal and distal TE cell populations display distinct gene expression patterns

Having determined that ALDH activity is a suitable criterion for enriching for stem/progenitor cells, we created mRNA-seq libraries for 34 samples (7 proximal EpCAM+/ALDH-, 9 proximal EpCAM+/ALDH+, 10 distal EpCAM+/ALDH-, 8 distal EpCAM+/ALDH+) from 12 generally healthy donors. Following data pre-processing (see Methods) we applied NGS checkmate (Lee et al., 2017) to verify that each library originated with the individual indicated by our records (Supplementary Figure 3). As a final quality check, we performed Gene Set Enrichment Analysis (GSEA) (Subramanian et al., 2005) using expressed ALDH family proteins and found that ALDH gene expression is significantly up-regulated in EpCAM+/ALDH+ samples (Supplementary Figure 4).

Principal component (PC) analysis has found the 4 cell populations segregate into visually distinct groups (Figure 3A). To determine how relevant the samples’ apparent segregation might be to identifying factors which may account for differences between the proximal and distal TE, we looked for significant correlations between variables of interest and potentially confounding factors. Of the potential covariates we tested, ALDH activity and region of origin correlated most strongly with PC1 and PC2. Importantly, the individual that donated sample material was not significantly correlated with any of the first PCs (Figure 2B).

**Figure 3.**
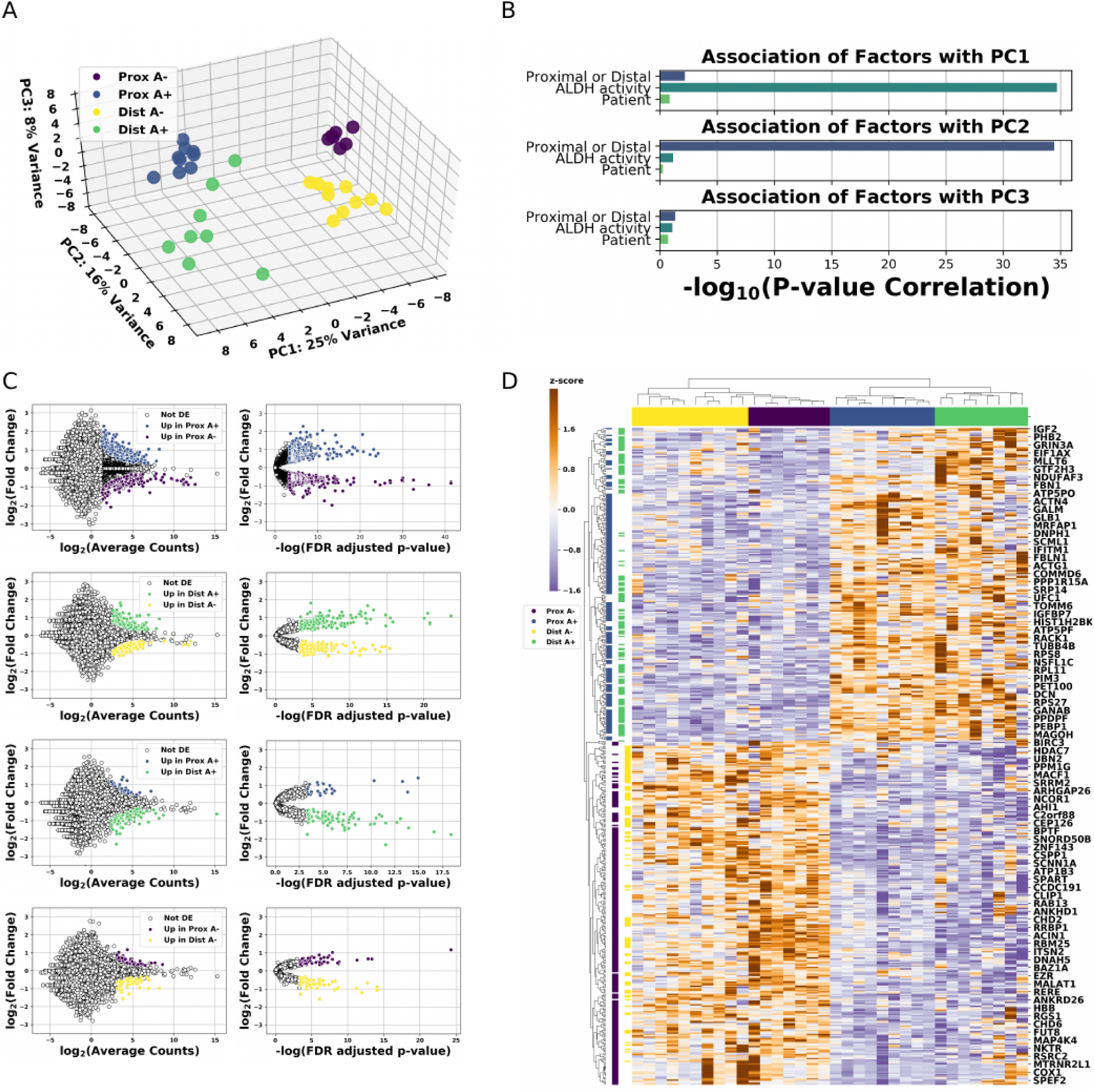
Gene expression data from primary human proximal and distal EpCAM+/ALDH- and EpCAM+/ALDH+ cell populations. (A) Principle component (PC) plot of the first three PCs from human TE mRNA-seq samples. Each dot belongs to a population indicated by the dot’s color. (B) Bar plots indicating the significance of the association between the indicated PC and the covariate listed on the y-axis. (C) Differentially expressed genes in human TE populations. Each MA and volcano plot pair displays genes which are differentially expressed between the cell populations indicated in the MA plot’s legend. (D) Heatmap summarizing sample segregation by, and expression of differentially expressed genes. Colors at the top of each column indicate which population the column belongs to. Each row corresponds to a gene found to be differentially expressed in at least one comparison. The colors along each row indicate which population(s) the gene in that row is differentially expressed in.

Having observed the high correlation between ALDH activity and region of sample origin with PC1 and PC2, we performed differential expression analysis using the DESeq2 (Love et al., 2014) R package. As expected, stem and differentiated cell enriched populations recover greater numbers of differentially expressed genes than comparisons between proximal and distal populations (Figure 3C). An overview of expression differences, which display expression trends distinguishing proximal and distal TE, is given in Figure 3D.

### Stem cell enriched populations exhibit increased Wnt signaling compared to differentiated cell enriched populations

To contextualize our differential expression results, we conducted gene ontology enrichment analysis using genes that are upregulated in EpCAM+/ALDH+ populations compared to their EpCAM+/ALDH- counterparts (Figure 4A) as well as GSEA (see Supplementary File 1 for gene sets). Stem/progenitor cells can play a role in malignant transformation and so we have begun by searching genes up regulated in EpCAM+/ALDH+ populations for enrichment in Disease Gene Network (Yu et al., 2015). We have found top hits relating to metastatic disease (Figure 4B). Querying GO Biological Process also recovered ‘cell-cell signaling by wnt’ as a prominent, statistically significant result (Figure 4C). We continued by performing GSEA on EpCAM+/ALDH+ vs. EpCAM+/ALDH- cell enriched populations. GSEA identified enrichment of the Hallmark Wnt/β-Catenin Signaling gene set (Figure 4D). β-Catenin and TCF family transcription factors are important mediators of canonical WNT signaling, which is an important pathway in maintaining SC self-renewal and cancer. We have found that distal EpCAM+/ALDH+ cell populations also show significant up regulation of *β-Catenin* (*CTNNB1*) and *TCF7* compared their EpCAM+/ALDH- counterparts. To see if WNT signaling may distinguish proximal from distal EpCAM+/ALDH+ populations, we examined the expression fold changes between genes annotated as involved in the WNT Signaling GO Biological Process. Expression of genes associated with WNT signaling is comparable between proximal and distal EpCAM+/ALDH+ populations (Figure 4G).

**Figure 4.**
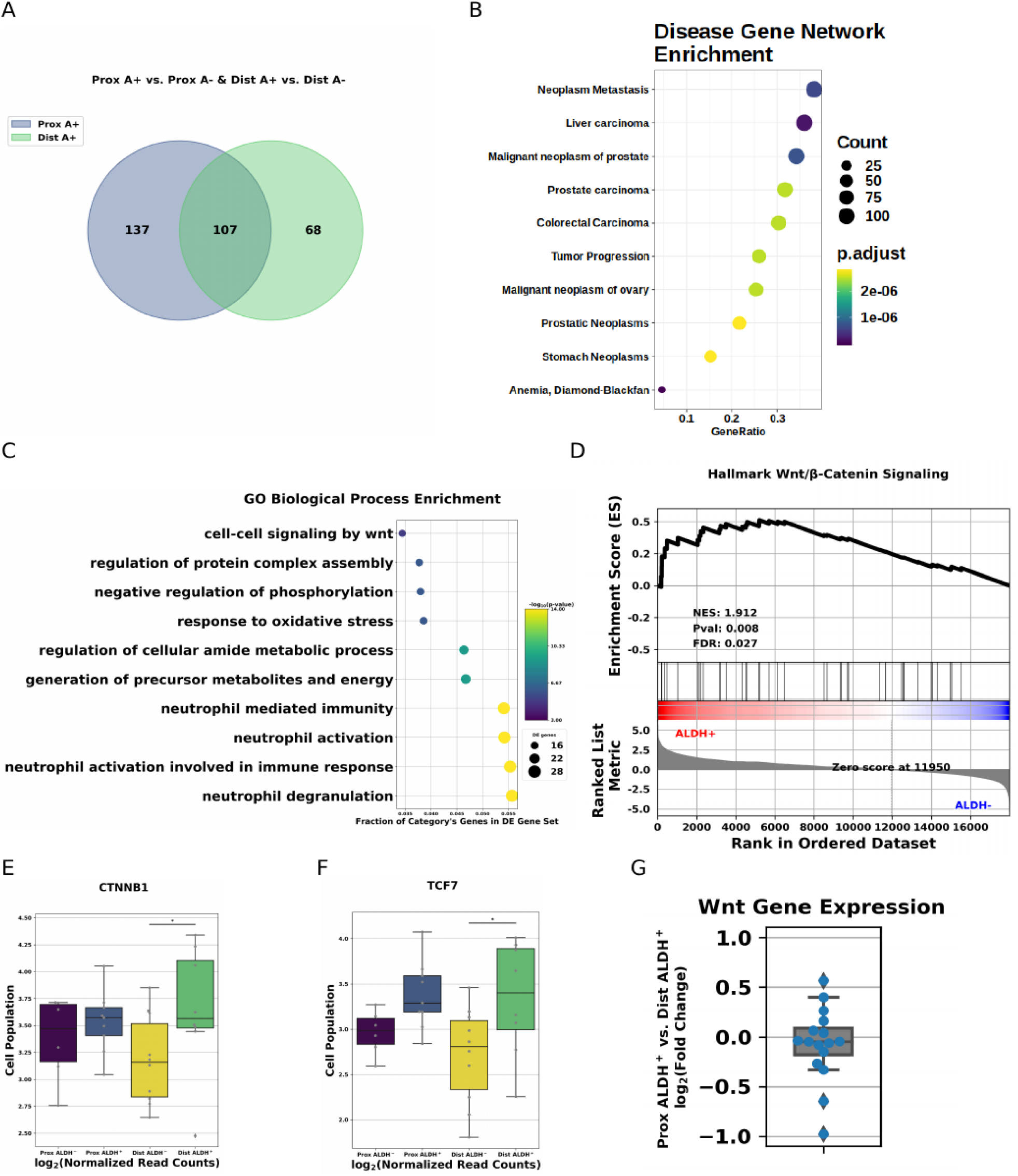
Functional significance of differentially expressed genes. (A) Venn diagram summarizing the extent to which differentially expressed genes from comparisons made in Figure 3C overlap. (B) Top 10 Disease Gene Net enrichment results sorted by multiple test corrected p-value for the 312 genes upregulated in proximal and distal EpCAM+/ALDH+ cell populations compared to their EpCAM+/ALDH- counterparts. (C) Top 10 GO Biological Process enrichment results sorted by hits/background gene for the same 312 genes as Figure 4B. (D) GSEA results for Hallmark Wnt/β-Catenin signaling. (E) Boxplots displaying β-Catenin signaling in each of the four cell populations. (F) Boxplots plots displaying TCF7 expression in each of the four cell populations. (G) Log_2_(Fold Change) for cell-cell signaling by Wnt genes between proximal and distal EpCAM+/ALDH+ cell populations.

### Inflammatory signaling is more pronounced in distal TE cell populations

The same gene set enrichment analysis that identified up-regulation of Wntβ-Catenin signaling in EpCAM+/ALDH+ compared to EpCAM+/ALDH- samples also found a pronounced up regulation of genes in the ‘Hallmark Inflammatory Response’ gene set in EpCAM+/ALDH- cells compared to EpCAM+/ALDH+ cells. As mentioned, inflammatory signaling is known or thought to contribute to malignant transformation in a variety of contexts. This led us to wonder how extensively activation of inflammatory pathways might differ between different TE populations. GSEA identifies significant enrichment of the Hallmark Inflammatory Response in differentiated cell populations from both the proximal and distal TE (Figure 5A-B). The distal TE EpCAM+/ALDH- samples displayed a greater enrichment for inflammatory signaling genes than did proximal EpCAM+/ALDH- (Figure 5A-B). Returning to the results of our differential expression analysis, we have noted that certain genes that are known to play roles in promoting or mediating pro-inflammatory responses are up-regulated in the distal TE, and that among these genes are *IGF2* and *TNFAIP2* (Figure 5C-D). Additionally, comparing all EpCAM+/ALDH+ samples to all EpCAM+/ALDH- samples showed an enrichment for Hallmark Inflammatory and TNF signaling in EpCAM+/ALDH- samples (Supplementary Figure 5).

**Figure 5.**
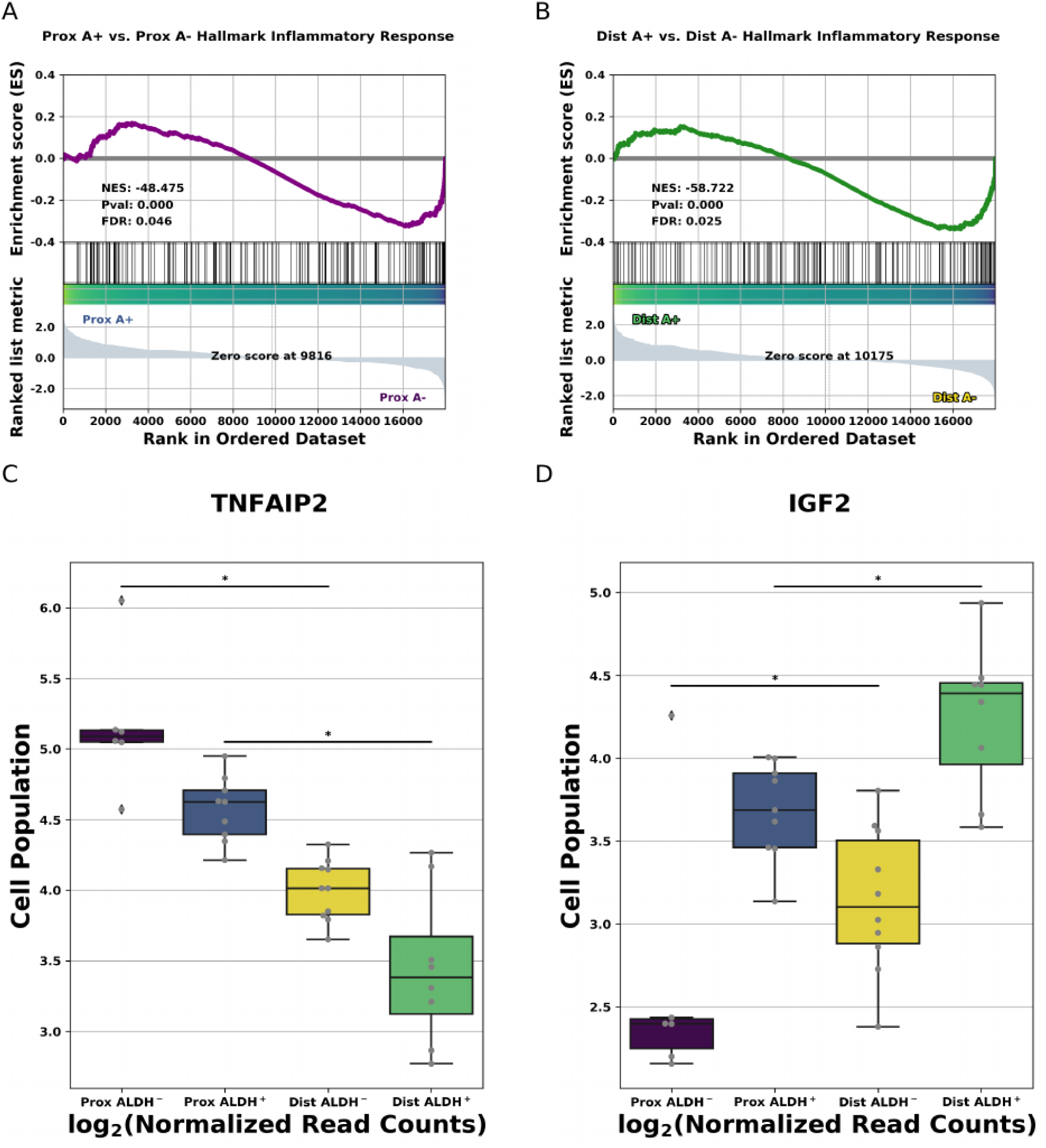
Inflammatory signaling in distal EpCAM+/ALDH- samples. (A) GSEA results for Hallmark Inflammatory Response in proximal EpCAM+/ALDH+ compared to proximal EpCAM+/ALDH- cell populations. (B) GSEA results for Hallmark Inflammatory Response in distal EpCAM+/ALDH+ compared to distal EpCAM+/ALDH- cell populations. (C) Boxplots displaying TNFAIP2 expression in each of the four cell populations. * indicated FDR adjusted p-value for indicated comparison is less than 0.1. (D) Boxplots exhibiting IGF2 expression in each of the four cell populations. * indicated FDR adjusted p-value for indicated comparison is less than 0.1.

To follow up on these observations we performed weighted gene co-expression network analysis (WGCNA, (Zhang and Horvath, 2005)). We observed 22 groups of genes displaying concerted changes in expression across the 4 conditions. We found one network (‘black’) particularly interesting due to its having the strongest correlation any cell type (Figure 6A). The genes comprising the ‘black’ module displayed a significant affinity for distal EpCAM+/ALDH- samples and a negative correlation with proximal EpCAM+/ALDH+ samples (Supplementary Figure 6). Pathway enrichment analysis has indicated that genes, which comprise this co-expression network, are somewhat enriched for NF-κB signaling, as well as cytokine and toll-like receptor signaling (Figure 6B). To follow up on these findings, we decided to perform an orthogonal enrichment analysis using Qiagen’s Ingenuity Pathway Analysis Tool (IPA). Consistent with our GSEA and GO enrichment analysis NF-κB signaling was up-regulated in distal EpCAM+ samples (Figure 6C).

**Figure 6.**
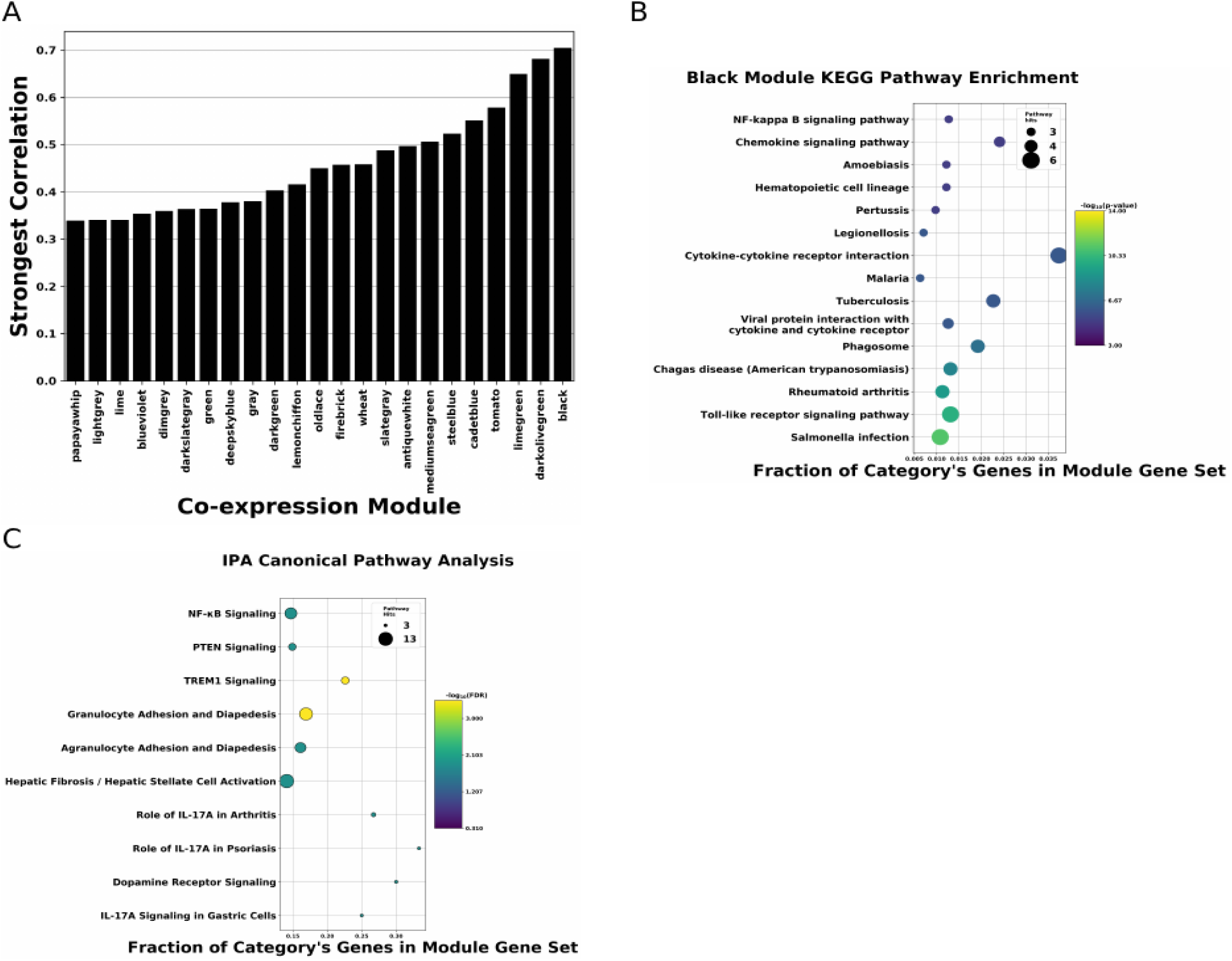
Distal TE gene co-expression networks. (A) Barplots indicating extent of correlation between any of the 4 cell types and a given co-expression module. Each position on the x-axis corresponds to a co-expression module. The height of the bar corresponds to the highest absolute value of correlation of that module with any of the 4 conditions. (B) KEGG Pathway enrichment results for ‘black’ co-expression network. (C) IPA heat map showing expression of pathways (names in column) in distal (left color-coded column) and proximal (right color coded column).

### Distal Fallopian tube epithelium is enriched for gene sets characteristic of HGSC

As has been mentioned, there is mounting evidence that a large fraction of HGSC originates in the distal region of the TE. HGSC encompasses at least four main molecular subtypes, but it is not clear if particular subtypes of HGSC are specifically associated with distal TE cell populations. Thus, we conducted differential expression analysis on each of the four main subtypes (1 vs. the other 3) for each molecular subtype using HGSC count data available from TCGA (Bell et al., 2011). GSEA finds each of these four gene sets was significantly up-regulated in the distal TE (Supplementary Figure 7A-D). Finding that the distal TE displays an upregulation of genes associated with HGSC, we wondered if distal TE ALDH+/EpCAM+ populations might express the same four gene sets more than distal TE ALDH-/EpCAM+ populations. Performing GSEA with the same four HGSC gene sets as above indicates that distal TE ALDH+/EpCAM+ populations tend to express the HGSC associated gene sets more highly, but only gene sets corresponding to the Immunoreactive and Proliferative HGSC subtypes have an FDR adjusted p-value below 0.05 (Figure 7A-D).

**Figure 7.**
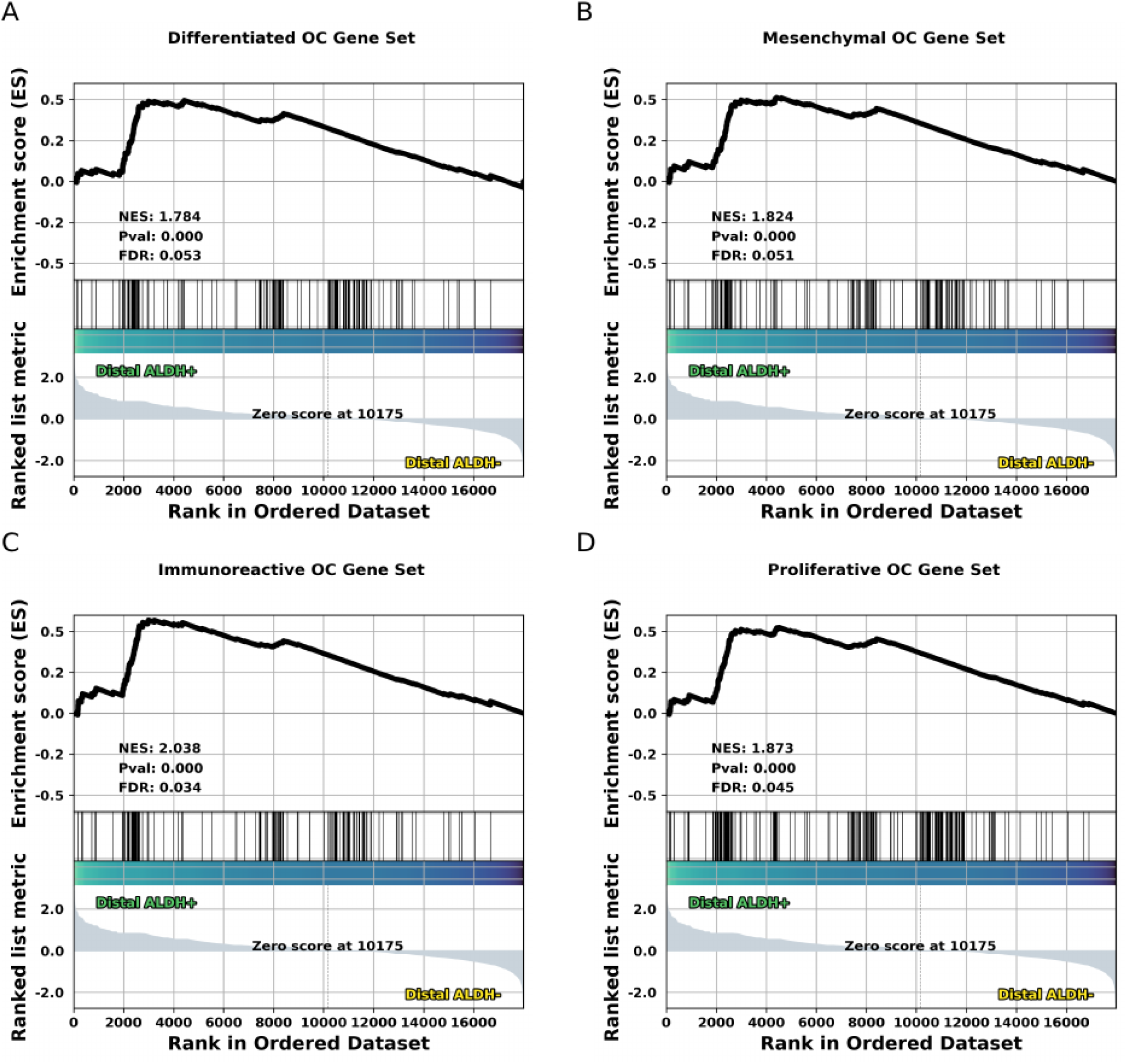
HGSC gene expression patterns in the human TE. (A-D) GSEA results for one of four gene sets corresponding to one of the 4 main molecular sub-types of HGSC identified by TCGA.

## Discussion

Quantitative organoid and genomic studies have been are largely absent with respect to the human Fallopian tube epithelium. Accordingly, understanding the reason for the distal TE’s susceptibility to malignant transformation is challenging. We observed a pronounced tendency towards organoid formation in distal compared to proximal bulk Fallopian tube patient samples. A cell population’s organoid formation tends to reflect the capacity for self-renewal and proliferation of the stem/progenitor cells within that population. Thus, differences in organoid formation between proximal and distal Fallopian tube samples are likely indicative of differences between the stem/progenitor cells of the proximal and distal regions of the Fallopian tube. Our bulk organoid formation results therefore strengthen the notion that the distal TE’s stem/progenitor cells or their environment differ in some way from those of the proximal region. These findings are consistent with observations that the distal region of the Fallopian tube more frequently contains putative HGSC precursor lesion (Kim et al., 2018; Lee et al., 2007; Schmoeckel et al., 2017).

Isolating stem/progenitor cells from more differentiated cells is a necessary pre-requisite for understanding cell lineage dynamics in a variety of contexts. Using ALDH activity assayed by FACS/AldeRed, we have observed EpCAM+/ALDH+ populations contribute a larger fraction of a given tissue sample’s organoids than the corresponding EpCAM+/ALDH- population. This leads us to conclude that ALDH activity is a reasonable heuristic for enriching TE cell isolates for putative stem/progenitor cells.

We set out to understand how proximal and distal TE populations differ, and how these differences may help explain the evident tendency of the distal TE towards malignant transformation. Gene ontology and gene set enrichment analysis data indicate EpCAM+/ALDH+ populations (which we take to be enriched for stem/progenitor cells) employ canonical Wnt/β-Catenin signaling more extensively than cells in the (generally more differentiated) EpCAM+/ALDH- populations. Our confidence in this conclusion is strengthened by the presence of *β-Catenin* and *TCF7* among the differentially expressed genes found between putative SC/progenitor and differentiated cell enriched populations. This conclusion is consistent with observations made by the Kessler group (Kessler et al., 2015). However, our observations of primary TE gene expression data did not find significant involvement of Notch signaling, which was previously identified as a requirement for maintaining long-term TE organoid cultures. This may indicate that human TE SCs rely on other mechanisms of Wnt signaling regulation *in vivo*. However, we cannot exclude the possibility that technical limitations inherent to our study obfuscated evidence of Notch signaling.

We observe a significant enrichment of inflammatory genes in differentiated cell enriched populations from both the distal and proximal regions. The increased expression of TNFAIP2 and IGF2 in the cells isolated from the proximal TE lends further credibility to the inflammatory hypothesis. IGF2 is present in follicular fluid and has recently been shown to promote malignant transformation in immortalized TE cell lines (Hsu et al., 2019). TNFAIP2 is thought to be involved in inflammatory angiogenesis and has been reported to down regulate NF-κB (Jia et al., 2018; Thair et al., 2015). If we suppose TNFAIP2 has a role in controlling NF-κB signaling in the TE, we would expect to see NF-κB signaling activity to be reduced in the proximal TE. This expectation is fulfilled by our weighted gene co-expression network analysis and orthogonal Ingenuity Pathway Analysis which both find increased NF-κB signaling in distal differentiated cell populations. NF-κB signaling is known to increase cellular proliferation and down-regulate TP53 mediated DNA damage response. Finding NF-κB signaling more active in primary human cell mRNA-seq data implicates NF-κB signaling in the distal TE’s evident susceptibility to malignant transformation and provides new, observational, evidence supporting the incessant inflammatory hypothesis. The pronounced inflammatory/NF-κB signaling Increased NF-κB signaling in differentiated cell population may lead to formation of altered niche increasing the propensity of stem cells to malignant transformation. However, we also cannot exclude that more differentiated cell population of distal TE may also to succumb to malignant transformation instead of less differentiated cell types.

The origins of HGSC are of considerable relevance to human health. We sought to assess gene expression patterns in primary human TE cell populations, to see if we might discern similarities between a particular molecular subtype of HGSC and a particular region of the TE. We find that the distal TE is significantly enriched for gene sets characteristic of HGSC subtypes. This is consistent with histological studies which find STICs occur more frequently in the distal region of the TE, though it does not help us determine for which subtype a given STIC is likely to give rise to.

While we are excited by these findings, we wish to stress some important limitations to our study. Though TNF family ligands are established regulators of NF-κB signaling, yet we do not observe significant differential expression of any TNF family genes. This may be addressed by analysis of stromally located immune cells, which may play in influencing the TE’s inflammatory response. A second peculiar finding is the absence of enrichment for cell cycle control genes. One would usually expect increased NF-κB signaling to be accompanied by a decreased DNA response activity and so eventual accumulation of mutations and genomic instability. We suspect our resolution is limited by the use of bulk mRNA-seq data, and the heterogeneity of epithelial cell populations in the TE. Future single cell studies will complement our current observations and garner important insight to HGSC’s pathogenesis and facilitate development of new approaches for its diagnosis, prevention and treatment.

## Materials and Methods

### Biosafety and ethical considerations

De-identified clinical samples were provided from the KU Cancer Center’s Biospecimen Repository Core Facility (BRCF) at KUMC along with relevant clinical information. Tissue specimens (Fallopian tubes - ampulla and fimbria) were obtained from women enrolled under the repository’s IRB approved protocol (HSC #5929) and following U.S. Common Rule. All patients provided written, informed consent in accordance with the BRCF IRB protocol. The samples were de-identified using OpenSpecimen. Samples were manipulated in a BSL II cabinet unless otherwise indicated.

### Collection and preservation of human TE samples

Proximal and distal Fallopian tube fragments were collected from standard surgical procedures for benign gynecological disease. The surgical tissue samples were collected on ice and examined by a surgical pathologist (maintaining the orientation and laterality of the sample), Only anatomically normal uterine tubes were used. Excised tissue fragments of 1 to 2 cm size were immersed in ice cold sterile phosphate-buffered saline (PBS; Corning, Corning, USA, catalogue #21-040-CM), washed once and transferred into 1 ml cryopreservation medium M-TE-N1. M-TE-N1 consists of 75% TE-M1, 15% fetal bovine serum (FBS; MilliporeSigma, Massachusetts, USA, catalogue #ES009-M), and 10% dimethyl sulfoxide (DMSO; MilliporeSigma, catalogue #D2650-100ML). TE-M1 consists of 5% FBS, 4 mM L-glutamine (Corning, catalogue #25-005-CI), 1 mM sodium pyruvate (Corning, catalogue #25-000-CI), 100 units ml^-1^ 100 ug ml ^-1^ penicillin/streptomycin (PS; Corning, catalogue #30-002-CI), 5 µg ml^-1^ insulin human (MilliporeSigma, catalogue #I9278),10 ng ml^-1^ epidermal growth factor human (EGF; MilliporeSigma, catalogue #E9644-.2MG), 10 ng ml^-1^ fibroblast growth factor-basic human (FGFb; MilliporeSigma, catalogue #F3133-10UG), 0.1 mM MEM non-essential amino acids (Corning, catalogue #25-025-CI), 0.1 mM MEM essential amino acids (Corning, catalogue #25-030-CI), 4% bovine serum albumin (BSA; MilliporeSigma catalogue #A3311-50G), and 500 ng ml^-1^ hydrocortisone (MilliporeSigma, catalogue #H0135-1MG) in 1:1 MCDB 105 (Cell Applications, San Diego, USA, catalogue #117-500) M199 (Corning, catalogue #10-060-CV) medium. Cryogenic vials with tissues were transferred to a minus 80°C freezer for short-term storage and shipped on dry ice to Cornell University.

### Bulk Fallopian tube organoid culture

Human distal and proximal tubal epithelial organoid culture was based on the previously established protocol (Kessler et al. 2013) with some modifications. Briefly, the snap-frozen tissues were thawed in the lab by incubating the tubes for 3 - 5 minutes at 37°C in the water bath. The thawed tissues were collected in 2D media [(Advanced DMEM/F12 (ThermoFisher, Waltham, USA, catalogue # 12634-028), supplemented with 5% FBS, 4 mM L-GlutaMax-I (ThermoFisher, catalogue #35050061), 10 µM Rho Kinase inhibitor Y-27632 (ROCKi; MilliporeSigma, catalogue #688000), 12 mM HEPES (ThermoFisher, catalogue # 15630-080), 10 µg ml^-1^ EGF human, 100 units ml^-1^ 100 ug ml^-1^ PS] and incubated for 3 minutes. The tissues were washed at least three times with PBS to remove extra blood and debris. The tissues were dissected into 2-mm sections and then minced finally using surgical blades. The tissue homogenate was incubated in 10 ml digesting buffer containing 0.5 mg ml^-1^ collagenase type I (ThermoFisher, catalogue #17100-017) in Advanced DMEM/F12, medium containing 12 mM HEPES and 3 mM CaCl_2_ for 45 min at 37°C in a water bath with rigorous shaking every 10 minutes. The tissues were minced again, centrifuged at 300xg at 22°C, pellets were suspended in 30 ml 2D media, passed through a 100, 70, and 40 µm cell strainer. The collected cell suspension was then spun at 300xg for 7 minutes at +4°C and the cell pellet was suspended in 3D media. The 3D media consisted of Advanced DMEM/F12, 10% R-Spondin1 conditioned medium, 25% Wnt3a conditioned medium, 4 mM L-GlutaMax-I, 12 mM HEPES, 1% N2 (ThermoFisher, catalogue # 17502048), 2% B27 (ThermoFisher, catalogue # 12587010), 100 ng ml^-1^ FGF10 human (Peprotech, Rocky Hill, USA, catalogue # 100-26-25ug), 1 mM Nicotinamide (MilliporeSigma, catalogue # N0636-100G), 10 µM ROCKi, 100 ng ml^-1^ Noggin human (Peprotech, catalogue #120-10C-20ug), 2.5 µM transforming growth factor-β Receptor inhibitor Kinase inhibitor VI (TGF-β Ri Ki VI; MilliporeSigma, catalogue #616464-5MG), 10 ng ml^-1^ EGF human, 1 µg ml^-1^ Osteopontin (OPN; MilliporeSigma, catalogue #120-35), 100 units ml^-1^ 100 ug ml^-1^ PS.

A total of 5 × 10^4^ cells were suspended in 3D media and mixed with growth factor reduced Phenol Red free Matrigel (Corning, catalogue #356231) in the ration of 30:70. This mixture was gently spread around the rim of a 12 well plate (rim assay). The plates were allowed to incubate for 20 minutes at 37°C in a 5% CO_2_ incubator. Once the Matrigel was solidified, 500 µl of 3D media was added to rim assays and incubated at 37°C. 10 µM p38 inhibitor (p38i; MilliporeSigma, catalogue # S7067-5MG) was added to the 3D media for the first 4 days and discontinued thereafter. The media was changed every second day and depending on the culture density the rim assay was passaged every 12 days.

Centrifugation was carried out at 4°C and 300xg unless otherwise noted. Statistical analysis on organoid data was performed with unpaired student *t*-test. All data represented in the figures with mean ± SD. A difference was considered statistically significant at a value of *P* < 0.05.

### Immunofluorescence analysis of organoids

Immunofluorescence analysis of paraformaldehyde-fixed paraffin embedded or frozen organoids was carried out using modified previously established (Flesken-Nikitin et al., 2016; O’Rourke et al., 2016).Briefly, at 22°C the culture medium from individual organoid rim assays was removed without disturbing the organoid/Matrigel rim mixture. The assay plate was placed on ice and 1 ml of ice cold Fixation buffer was added for 3.5 hours. Fixation buffer consists of 4% paraformaldehyde in 1x PME buffer. A 10x PME buffer consists of 500 mM 1, 4-Piperazinediethanesulfonic acid (PIPES; Bioworld, Dublin, USA, catalogue #41620140-1), 25 mM Magnesium Chloride, and 0.5 M Ethylenediaminetetraacetic acid (EDTA; MilliporeSigma, catalogue #AM9260G). After the fixation, continue the work at 22°C. The Fixation buffer was taken out from the middle of the wells, followed by addition of 0.5 ml PBS supplemented with 0.2% Triton X 100 (MilliporeSigma, catalogue #T8787-50ML and 0.05% Tween (MilliporeSigma, catalogue #T2700-100ML). The organoid suspension was collected in a 1.7 ml centrifuge tube. Wide bore yellow tips were used from this point. Organoids were centrifuged at 300xg at 4°C, washed three times with PBS 0.2% Triton X 100 0.05% Tween, and once with PBS. The organoid pellet was suspended for dehydration in 600 µl 70% ethanol and incubated overnight at 22°C. The next day the organoid pellet was dissolved (with taking out as much 70% ethanol as possible) in 50 µl of melted Histogel (ThermoFisher, catalogue #17985-50). The suspension forming a droplet was pipetted on a Parafilm lined petri dish and solidified at 4°C for 10 minutes. The solidified Histogel droplet containing organoids was stored in 70% ethanol and later processed for paraffin embedding. The organoids were sectioned 4 µm thick and subjected to immunofluorescence staining using xylene deparaffinization and serial rehydration over a graded ethanol series. The antigen retrieval was performed using 10 mM sodium citrate buffer at pH 6.0 for 10 minutes. The primary antibodies against PAX8 (Abcam, Cambridge, UK, catalogue #ab189249), ALDH1A1 (Abcam, Cambridge, UK, catalogue #ab52492), and ACTUB (Sigma, St. Louis, USA, catalogue #T6793) were incubated in a humidified chamber overnight, followed by incubation with secondary antibodies (Donkey anti-Rabbit IgG (H+L) and Donkey anti-Mouse IgG (H+L) Alexa Fluor 488) for 1 hour at room temperature. Sections with no primary antibody served as negative controls. The stained sections were mounted in ProLong Diamond Antifade Mountant with DAPI reagent (Thermo Fisher). Confocal images were acquired using a Zeiss LSM 710 confocal microscope through the Cornell University Biotechnology Resource Center. The image data was merged and displayed with the ZEN software (Zeiss).

### Preparation and collection of human TE FACS samples

Human Fallopian tube samples were removed from liquid nitrogen and thawed at 37°C for 3 minutes before being removed from the cryo-preservation vial and being rinsed 3 times with 15ml 1x PBS. Each sample was then dissected and minced to reveal as much of the mucosa as possible, any coagulated blood was scraped away. Samples were then incubated at 37°C for 45 minutes in Digestion Buffer, shaking every 10 minutes. Samples were then collected by centrifugation, placed in 2D Culture media and mechanically dissociated using a 5 ml serological pipette. Sample fragments were then ground with a mortar and pestle using a 300 µm filter before being further dissociated with 5 strokes of a loose Wheaton Dounce homogenizer. Samples were successively filtered through 100, 70, and 40 µm mesh filters before being collected by centrifugation and being re-suspended in 2D FACS media [Advanced DMEM/F12, supplemented with 1% N2, 2% B27, 1 mM Nicotinamid, 1 mM N-Acetyl-L-Cysteine (MilliporeSigma, catalogue # A9165-25G), 10 µM ROCKi, and 100 units ml^-1^ 100 ug ml^-1^ PS]. Samples were successively filtered through 100, 70, and 40 µm mesh filters before being collected by centrifugation and being re-suspended in 2D FACS media. For detection of ALDH enzymatic activity, sample cells were suspended in the AldeRed Assay Buffer and processed for staining with the AldeRed ALDH Detection Assay (MilliporeSigma, catalogue #SCR150) according to the manufacturer’s protocol. At this point roughly 250,000 cells were set aside for the Diethylaminobenzaldehyde (DEAB, ALDH inhibitor), EpCAM, and compensation controls. DEAB control was prepared according to manufacturer’s instructions as well. Samples/isotype control were stained with EpCAM/conjugated isotype for 1 hour at 5°C according to manufacturer’s instructions. Appropriate sample suspensions were stained with SYTOX Blue prior to sorting on a BD FACS Aria III using 450/50, 610/20, and 696/40. Sorted cells were collected directly into 750 µl Trizol-LS (Fisher Scientific) as described (Jadhav et al., 2017).

### ALDH activity segregated organoid culture

Approximately 3×10^5^ viable EpCAM+/ALDH+ and EpCAM+/ALDH- collected by FACS into FACS media (described above) after the preparation described above. Collected cells were recovered by centrifugation at 300xg for 15 min at 4°C. Most of the remaining liquid was decanted, and roughly 50x the remaining volume in Matrigel was added to the sample and gently mixed by pipetting. 20-30 µl droplets were then plated and allowed to sit for 30-40 minutes before the addition of 250 μl T-media. Media was changed ever two days. Cultures were passaged every week. Passaging was done by dissociating the organoids by pipetting in ice-cold 3D media containing p38i. Organoid cultures were then re-plated as described above, and dissociation to single cells was verified using bright field microscopy.

### mRNA-seq library preparation and data pre-processing

3’prime mRNA-seq libraries containing unique molecular identifiers (UMIs) were prepared using Lexogen’s QuantSeq Kit (Lexogen, Vienna, Austria, catalogue # 015.24, #081.96) according to the low-input protocol. Optimal barcodes were assigned to each sample by Lexogen’s Index Balance Checker webtool (https://www.lexogen.com/support-tools/index-balance-checker/). Libraries were pooled and sequenced on an Illumina NextSeq 500 after undergoing QC by Agilient Fragment Analyzer.

De-multiplexed FASTQ files were inspected for quality using FASTQC (Brown et al., 2017). Reads were aligned to GRCh38 using the STAR two-pass method (Dobin et al., 2013). UMI-tools (Smith et al., 2017) was then applied to remove duplicate reads based on their UMI. Quality score and base re-calibration were then performed according to the Genome Analysis Toolkit best practices for mRNA-seq version 3.7. Sample identity was then verified using NGSCheckMate (Lee et al., 2017).

### Bioinformatics analysis

A raw read count matrix was generated using the featureCounts function of the Rsubread R package (Liao et al., 2019). Background and technical noise were reduced using the RUV-seq R package (Risso et al., 2014) before analysis differential expression analysis. Read count normalization and gene differential expression calls were made primarily with DESeq2 (Love et al., 2014) Gene and Disease Ontology (GO and DO) enrichment analysis was carried out using the clusterProfiler and DOSE R packages (Yu et al., 2012; Yu et al., 2015). Gene Set Enrichment Analysis (GSEA) was performed using the GSEApy python package (Chen et al., 2013; Subramanian et al., 2005). Weighted Gene Co-expression Network Analysis (WGCNA) was performed using the Weighted Gene Co-expression Network Analysis R package (Langfelder and Horvath, 2008; Zhang and Horvath, 2005). Ingenuity pathway analysis was done using the 1,000 most divergently expressed genes between all proximal EpCAM+ samples and all distal EpCAM+ samples using ‘epithelial pathways’ as a background set. For TCGA OV Analysis Gene set’s typifying the four main molecular sub-types of HGSC were obtained from TCGA array data (Network et al., 2011) and supplemented using TCGA mRNA-seq data obtained using the TCGAbiolinks R package (Colaprico et al., 2016).

## Acknowledgments

We would like to thank all members of the Nikitin Lab for their advice and support. This work has been supported by NIH (CA096823 and CA197160) and NYSTEM (C023050 and C028125) grants to AYN, the NIGMS (P20 GM130423), the KU Cancer Center’s Cancer Center Support Grant (P30 CA168524), and the Honorable Tina Brozman Foundation, Inc. to AKG, and fellowship funding from the predoctoral fellowships to IMR (NIH T32HD057854 and NYSTEM C30293GG) and postdoctoral fellowship to MB (NYSTEM C30293GG). The work on the Zeiss LSM 710 Confocal microscope was made possible by NIH grant S10RR025502 to the Cornell University BRC Imaging Facility. AKG is the Chancellors Distinguished Chair in Biomedical Sciences.

## Author contributions

IMR, AFN and AYN designed experiments. IMR, MB, and AFN performed experiments, IMR carried out bioinformatics analyses, AW and AKG led the collection of human materials and clinical information, IMR, AFN and AYN wrote the paper, and all reviewed the manuscript.

