## Supplementary Materials for "WNT and inflammatory signaling distinguish human Fallopian tube epithelial cell populations"

A

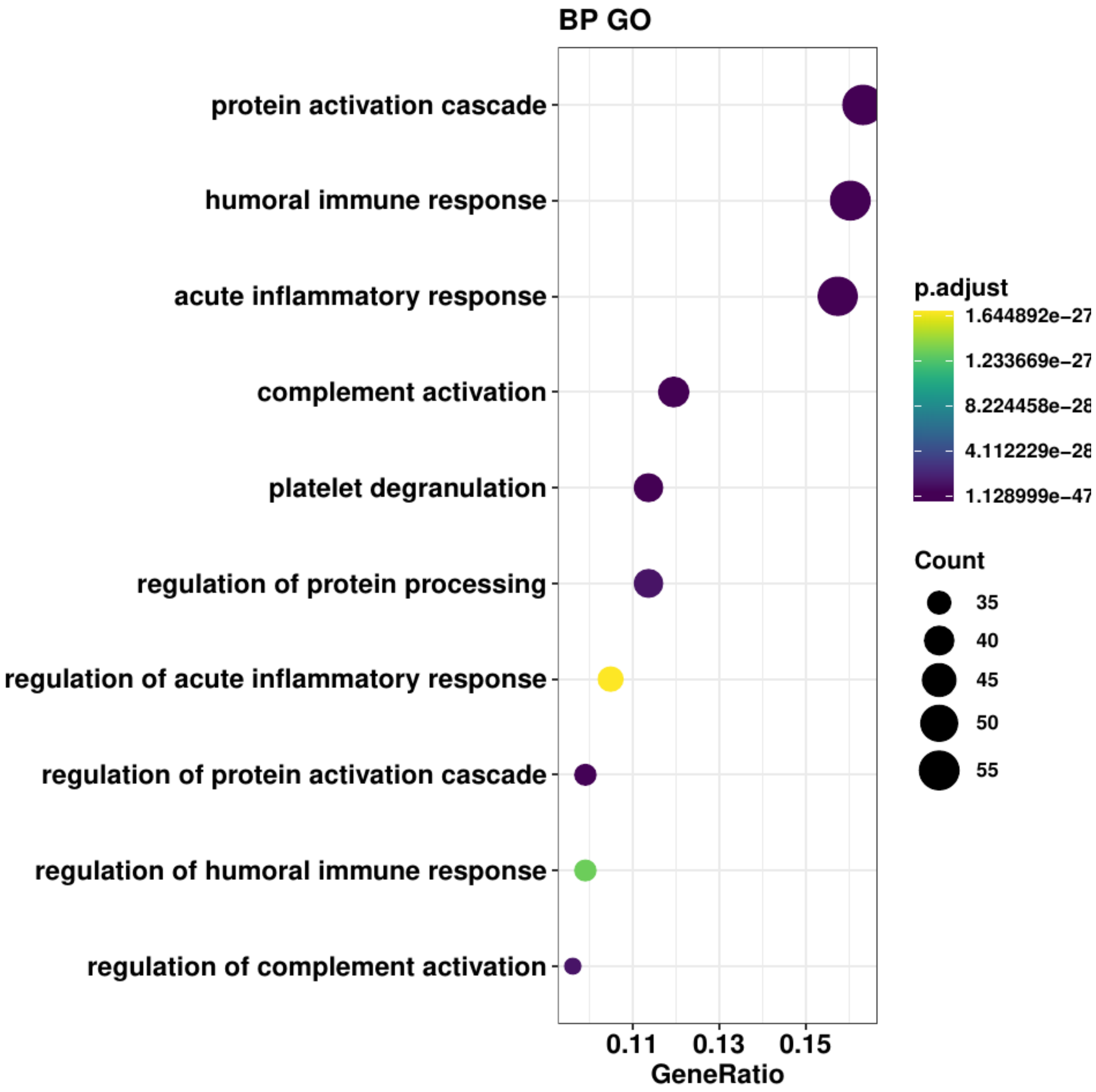


**Supplementary Figure 1. Follicular fluid protein ontology enrichment.** Dotplot indicating enriched GO biological process terms for proteins found in follicular fluid mass spectroscopy data (Lewandowska *et al.* 2019).

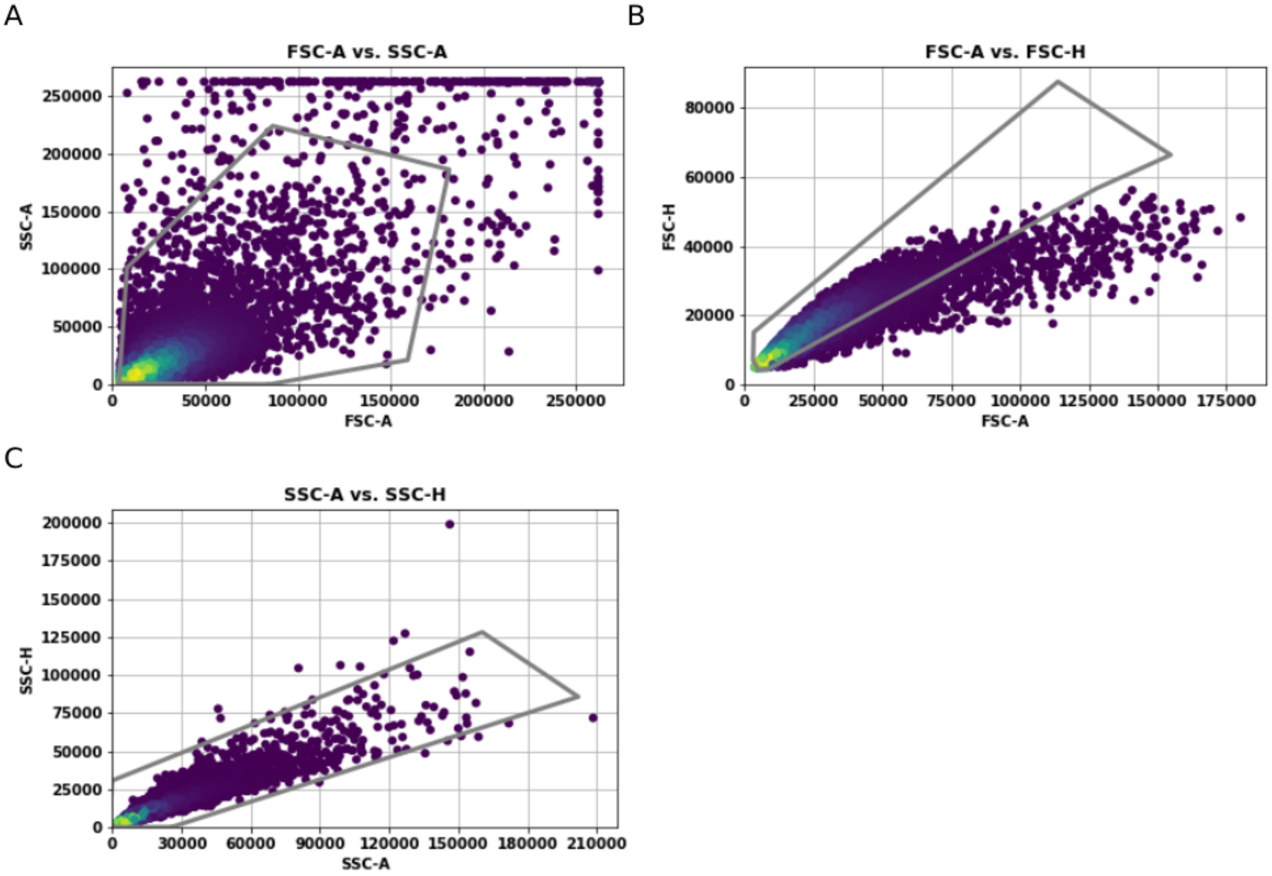


**Supplementary Figure 2. Representatives FACS light scatter gating.** (A) Forward vs. side scatter gate data. Points inside the grey polygone were interrogated further. (B) Forward scatter area vs. forward scatter height gate data. Points inside the grey polygone were interrogated further. (C) Side scatter are vs. side scatter height gate data. Points inside the grey polygone were interrogated further.

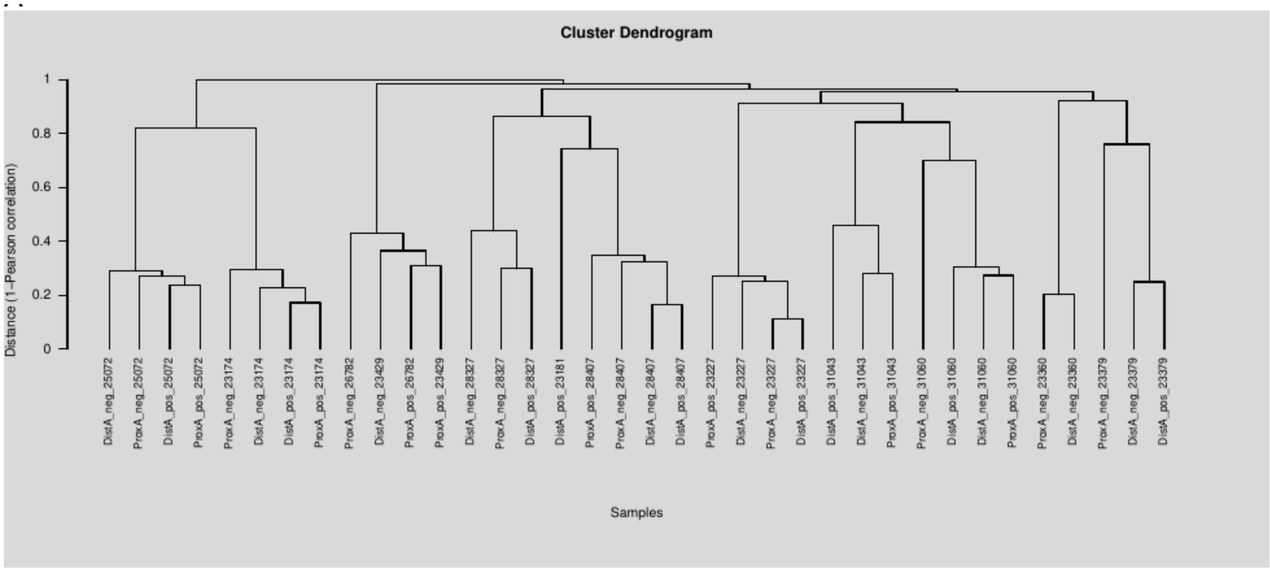


**Supplementary Figure 3. NGS Checkmate dendrogram.** SNP correlation matrix is used to cluster mRNA-seq samples by individual that donated material.

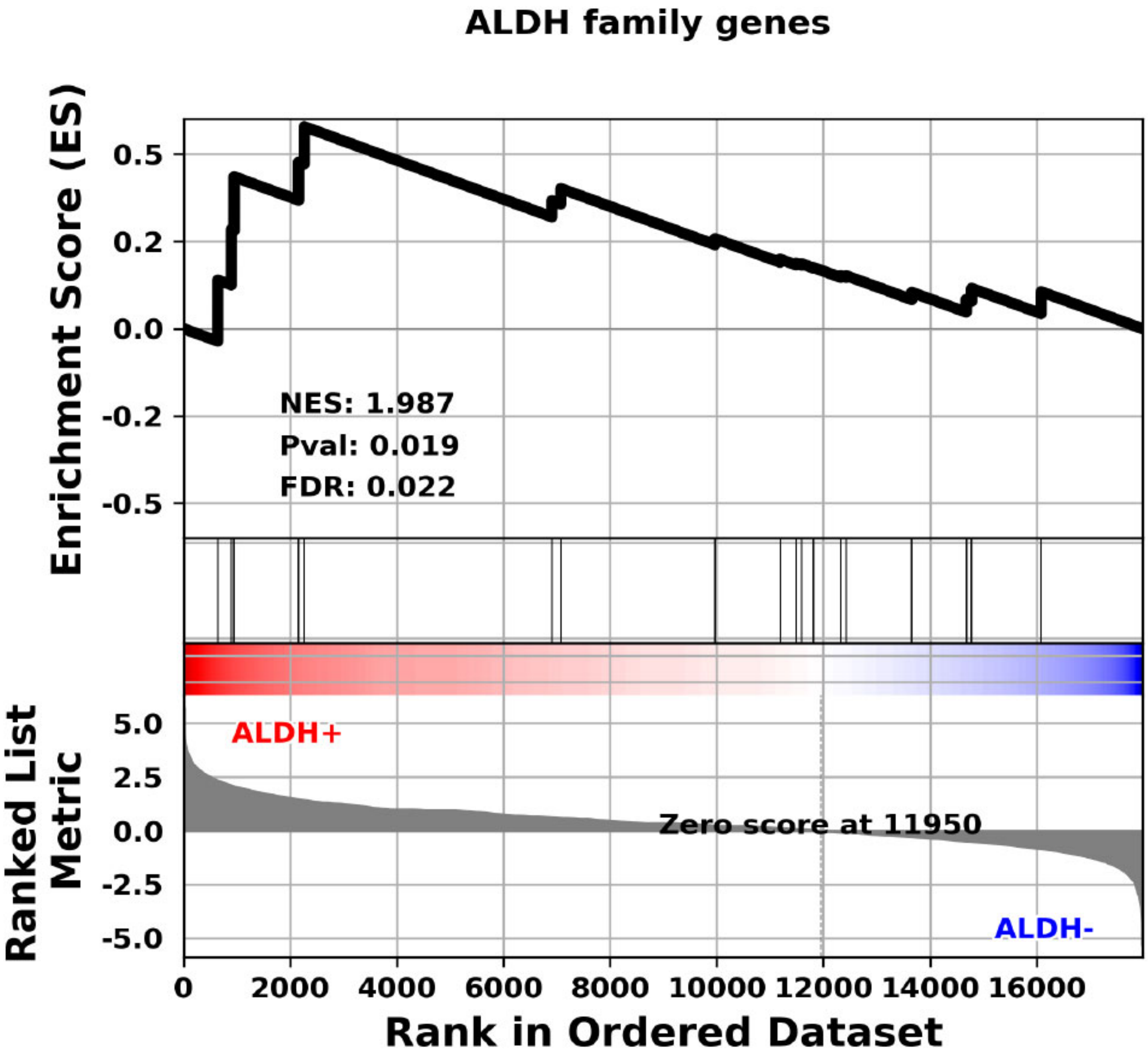


**Supplementary Figure 4. ALDH family GSEA.** GSEA using all ALDH family genes, which are present in primary human TE mRNA-seq data as a gene set.

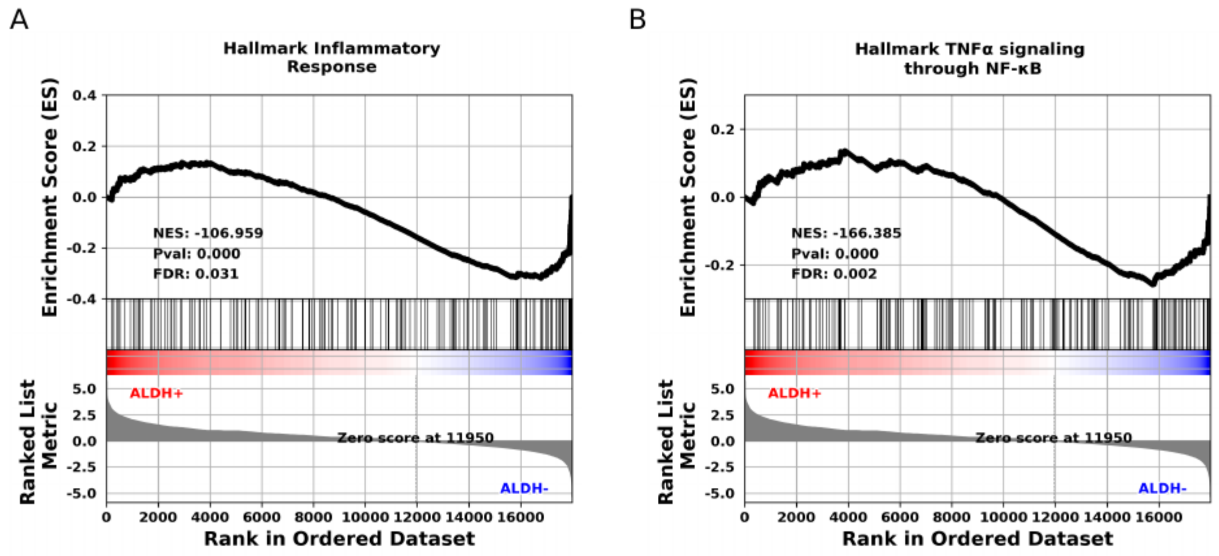


**Supplementary Figure 5. Two most statistically significant Hallmark gene set enrichment results.** (A) GSEA plot showing results for Hallmark Inflammatory Response gene set in all EpCAM+/ALDH+ samples compared to all EpCAM+/ALDH- samples. (B) GSEA plot showing results for Hallmark TNFα signaling through NF-κB gene set in all EpCAM+/ALDH+ samples compared to all EpCAM+/ALDH- samples.

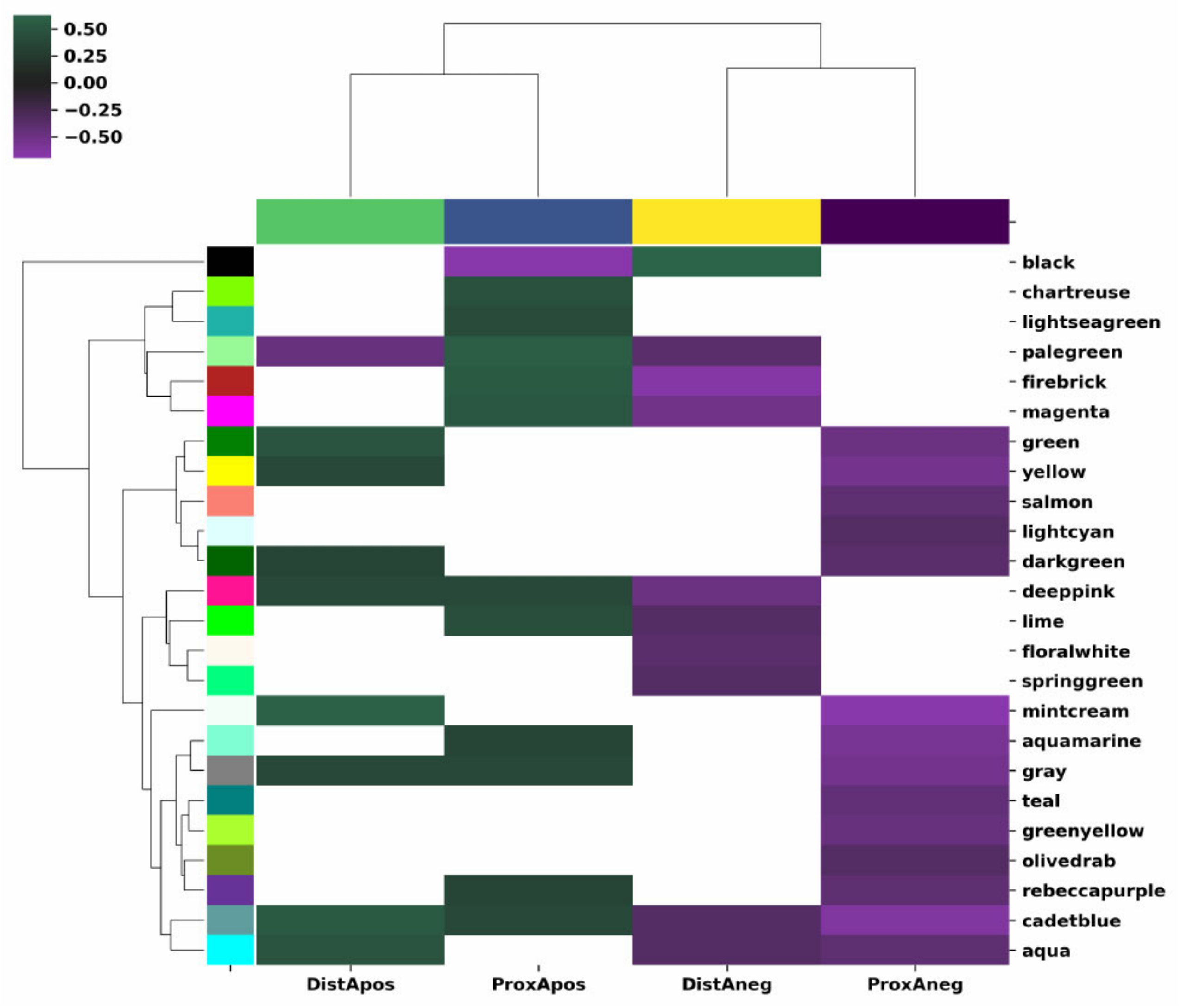


**Supplementary Figure 6. Weighted Gene Co-Expression Network analysis.** Each row corresponds to a gene expression module identified using WGCNA. Each column corresponds to one of the four cell types. Every colored (non-white) cell corresponds to a statistically significant ($p<0.05$) association between a co-expression module and a cell type. Green cells indicate a positive association between the co-expression module and the cell type. Purple cells indicate a negative correlation between the co-expression module and the cell type.

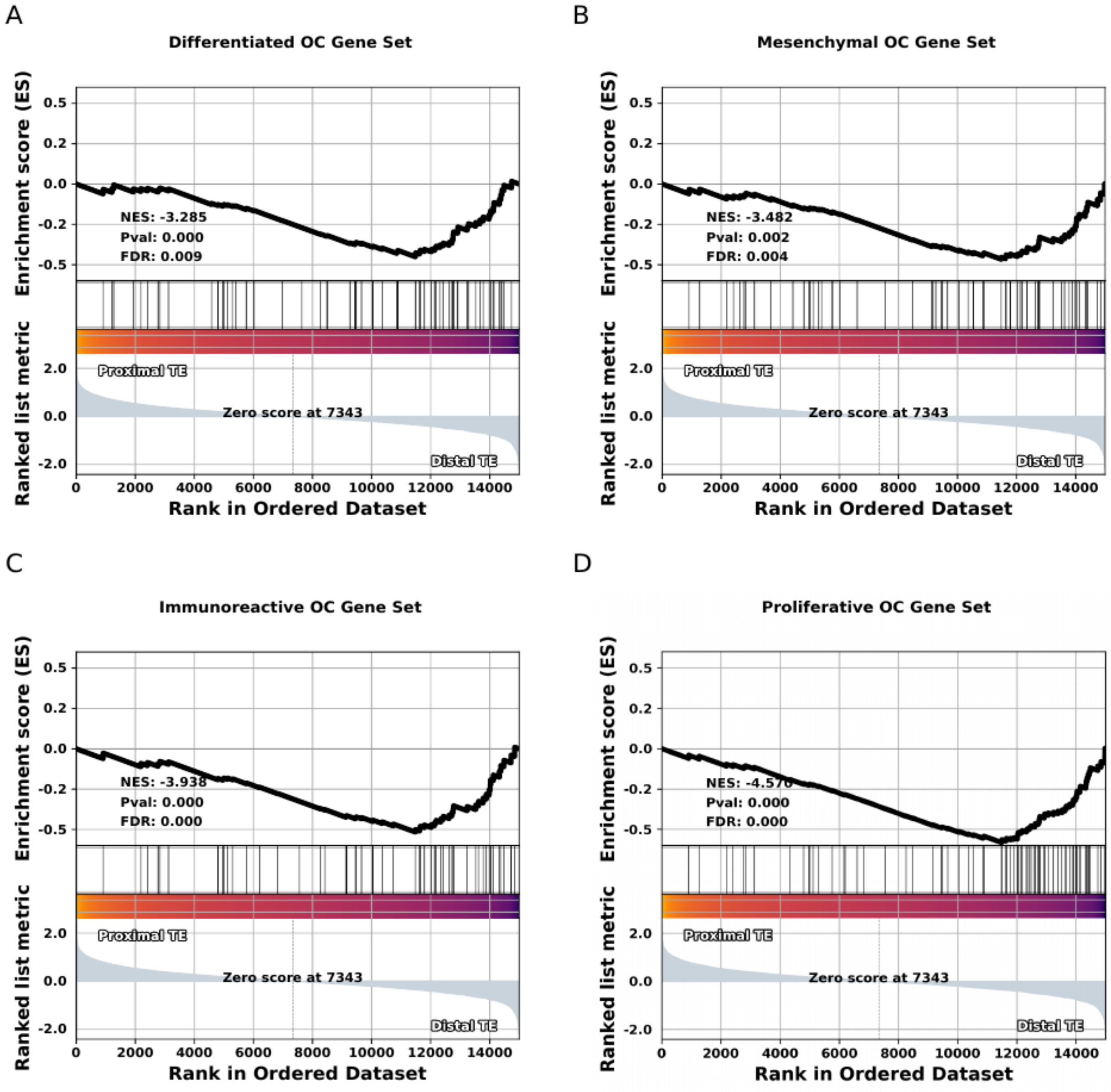


**Supplementary Figure 7. HGSC gene expression patterns in the proximal vs distal human TE.** (A-D) GSEA results for one of four gene sets corresponding to one of the 4 main molecular sub-types of HGSC identified by TCGA. All enrichment results shown pertain to all proximal region samples compared to all distal region samples.
